# Evolution of Boldness and Exploratory Behavior in Giant Mice from Gough Island

**DOI:** 10.1101/2020.09.10.292185

**Authors:** Jered A. Stratton, Mark J. Nolte, Bret A. Payseur

## Abstract

Island populations are hallmarks of extreme phenotypic evolution. Radical changes in resource availability and predation risk accompanying island colonization drive changes in behavior, which Darwin likened to tameness in domesticated animals. Although many examples of animal boldness are found on islands, the heritability of observed behaviors, a requirement for evolution, remains largely unknown. To fill this gap, we profiled anxiety and exploration in island and mainland inbred strains of house mice raised in a common laboratory environment. The island strain was descended from mice on Gough Island, the largest wild house mice on record. Experiments utilizing open environments across two ages showed that Gough Island mice are bolder and more exploratory, even when a shelter is provided. Concurrently, Gough Island mice retain an avoidance response to predator urine. F1 offspring from crosses between these two strains behave more similarly to the mainland strain for most traits, suggesting recessive mutations contributed to behavioral evolution on the island. Our results provide a rare example of novel, inherited behaviors in an island population and demonstrate that behavioral evolution can be specific to different forms of perceived danger. Our discoveries pave the way for a genetic understanding of how island populations evolve unusual behaviors.

## Introduction

Organisms that colonize islands experience unique environmental challenges that require novel solutions [1]. New kinds and distributions of resources on islands can select for new foraging strategies [2-4]. For many island colonizers, a loss of predators alleviates the need for behavioral or morphological defenses [2, 5-6]. If resources are difficult to acquire and predatory risk is absent, the optimal foraging-risk ratio is highly skewed, favoring exploration and boldness [7-9]. Island populations, therefore, provide opportunities to test hypotheses about the evolution of behavior in novel environments.

Populations that spread to islands often display behavioral changes. Many populations lose anti-predator behaviors [6]. Some island inhabitants are more easily approached by humans, a phenomenon Darwin called “island tameness” [10-11]. In some island rodents, territoriality and conspecific aggression are reduced compared to mainland populations [12-13]. Behavioral responses to island environments sometimes occur in the context of other phenotypic shifts. New behaviors are often accompanied by transitions in life history and morphology [12,14-15] in a pattern referred to as the “island syndrome” [16].

Although behavior is suspected to be an important component of adaptation to island conditions, the question of whether island populations harbor heritable changes in behavior has rarely been answered [13,17]. Demonstrating that behavior has evolved on islands requires evidence of genetic change in behavioral traits.

A promising organismal system for testing hypotheses about behavioral evolution in response to island environments is found on Gough Island, a remote volcanic island in the middle of the South Atlantic Ocean. Despite the remote location of Gough Island, over 2,700 miles from the nearest mainland, house mice (*Mus musculus domesticus*) colonized the island, probably via sealing ships from western Europe a few hundred generations ago [18-20]. In a remarkable phenotypic transformation, these mice evolved a body size twice that of their mainland counterparts [21-22]. Laboratory-born offspring of Gough Island mice (hereafter “GI mice”) maintain their unusual size, confirming that this morphological distinction has a genetic basis [23-24]. On Gough Island, mice live without the predators and without the human commensals they commonly exploit for food and shelter on the mainland [21,25]. The diet of GI mice is highly variable and seasonal, with invertebrates (mainly earthworms) and seeds being the most stable food source [22]. During the winter season, GI mice predate on endangered seabird populations that use the island as a nesting ground, leading to the deaths of an estimated 2 million chicks and/or eggs per year [26]. The combination of increased body size, novel consumption of birds, loss of predatory danger, and removal of human commensals predicts the evolution of increased exploration and boldness in GI mice. Because GI mice are western European house mice (*Mus musculus domesticus*), the same subspecies as the laboratory mouse [20], established methods in biomedical research can be applied to profile behavioral evolution.

In this article, we use GI mice to examine recent behavioral evolution on an island. By exposing juvenile and adult mice to novel environments with different levels of perceived risk, we uncovered multiple lines of evidence that GI mice are bolder (i.e. less anxious) and more exploratory than mice from a mainland reference strain. The detection of these differences in inbred strains raised in a common environment demonstrates that they are inherited. Our findings indicate that GI mice evolved enhanced boldness and exploration over a short timescale and in concert with other substantial phenotypic changes. This work lays the foundation for identifying the genetic changes responsible for behavioral evolution in organisms that colonize islands.

## Materials and Methods

### Mouse Strains and Husbandry

Inbred strains of GI mice and mainland mice were used throughout this study. In 2009, GI mice were live-caught and shipped to the University of Wisconsin School of Veterinary Medicine Charmany Instructional Facility, where a breeding colony was established [23]. GI mice for this study belonged to a strain maintained for 21-23 generations of brother-sister mating. At this stage of inbreeding, we expect most of the genome to be homozygous, allowing us to treat individual mice as replicates of the same genetic background. Mainland mice belonged to the WSB/EiJ inbred strain founded from breeding pairs caught in Maryland (purchased from the Jackson Laboratory, Bar Harbor, ME) and were maintained in the same colony as GI mice for the same time period. F1s were generated by crossing mice from the GI strain and the mainland strain in both maternal directions. All mice were housed in micro-isolator cages with corn cob substrate (1/8th inch; The Andersons Lab Bedding), with *ad libitum* access to food (Envigo 2020X Teklad Global Diet) and water. Breeders were provided with a higher fat chow (Envigo 2019 Teklad Global Diet) and a red mouse igloo (Bio Serv). All cages were provided with nesting material and irradiated sunflower seeds (Envigo). Cage changes occurred every 6-8 days or, in the case of a new litter, 10 days after parturition. The colony was kept in a temperature-controlled room (68-72°F) under a 12-hour light/dark cycle.

Mice used for behavioral testing were weaned 20-21 days after parturition and housed with one littermate of the same sex to reduce behavioral effects of within-cage hierarchies [27]. All mice were weighed to the nearest tenth of a gram during cage changeouts and after the last behavioral assay (*i*.*e*. weekly from ages 4-10 weeks old +/- 1 day).

### Behavioral Assays

All behavioral assays were conducted in a room separate from the main colony. Each subject was tested four times with the following regimen (see Figure 1A): open field (4 and 8 weeks old +/- 1 day), light/dark (9 weeks old +/- 1 day), and predator cue (10 weeks old +/- 1 day). All tests were conducted during the light phase of the light/dark cycle. Females were scored for stage of estrous cycle according to Caligioni [28] on the same day as testing, beginning with the second open field test. Twelve females (all from the mainland strain) were in estrous the same day as testing. These animals fell within the range of values for each trait measured in that strain and were included in the final analysis. The order of subjects tested within litters each day was randomized. Each test began by bringing the subjects’ cage into the room for a 30-minute acclimation period. The experimenter remained in the room out of sight of the subjects throughout the acclimation and testing periods except for transfer to and from the arena. The relevant arena for each test (see Figure 1B for examples) was placed in the center of the room next to a movable cart where a computer and video recording hardware were located. The arena was cleaned with 70% ethanol after each test and at least five minutes was allowed for the ethanol to evaporate before testing a new subject. All tests were video recorded using Debut Video Capture Software v 5.33 at default settings with Focus set to 0. A Logitech HD Pro Webcam C920 was positioned directly above the center of the arena. Two videos were taken before each test to assist in video analysis: an “empty” video containing a short recording of the empty arena, and a calibration video containing a short recording of the arena with specific positions marked. The “empty” video was used in aligning all images of the test recording. The calibration video was used in defining coordinates of regions of interest and converted pixels to millimeters. Subsequent subsections provide additional details of experimental design for each test.

**Figure 1:**
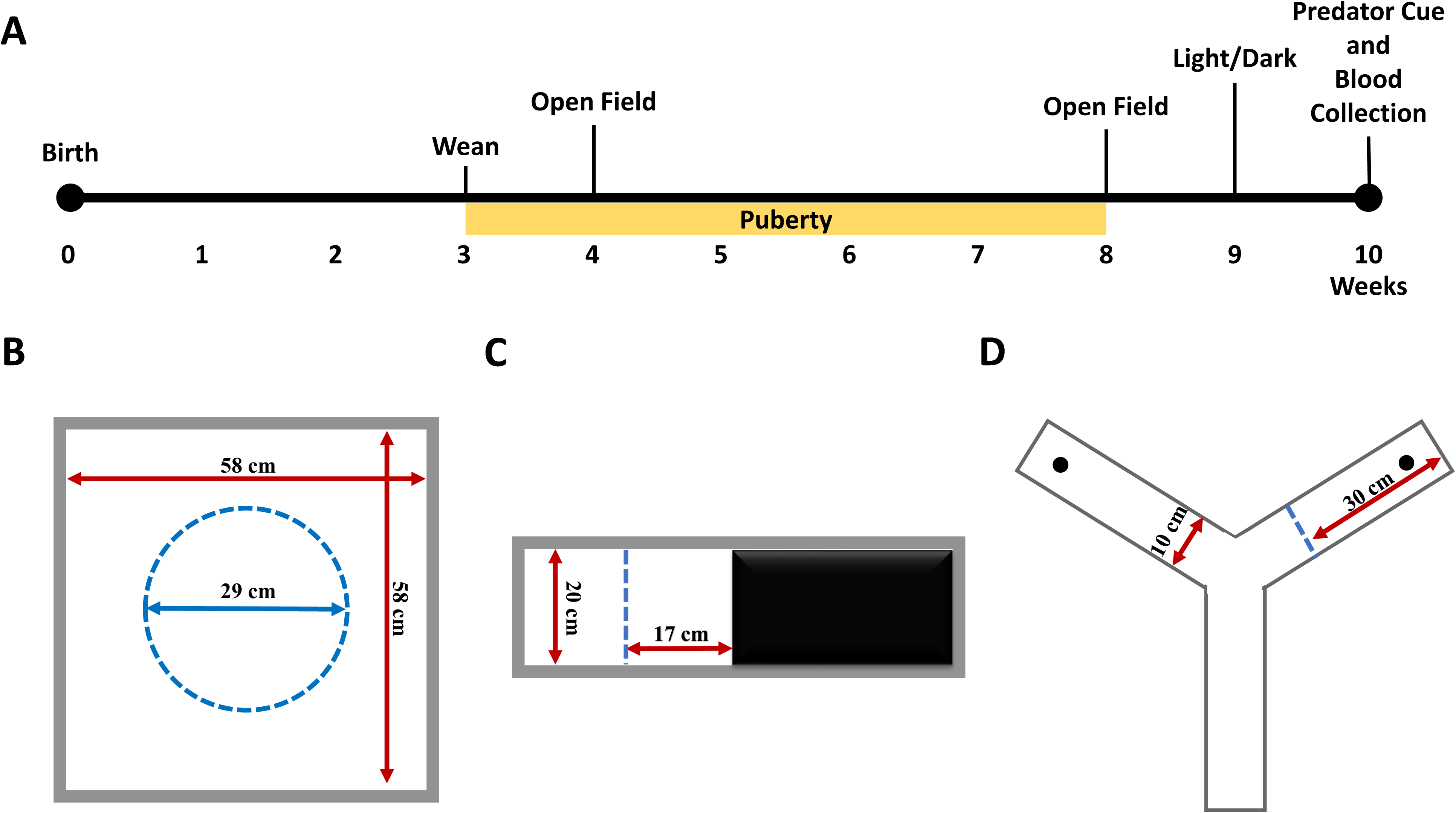
Schematic of testing design and arenas. A) Timeline of a subject mouse’s life. B) Schematic of the open field arena. The center defined during video analysis is outlined by the dashed circle. C) Schematic of the light/dark box. The subject has free access to both equally-sized chambers during the test. Ditance ot the center threshold is noted by the dashed line. D) Schematic of the y-maze use din the predator cue test. Each arm is equally sized and freely accessible. Threshold for each arm defined during video analysis is indicated in the right arm by the dashed line. Locations of the predator and control cues are noted with black dots.

### Open Field Test

The open field (58cm W x 58cm D x 58cm H) was constructed from expanded, white PVC (Grainger Industrial Supply). Lighting in the room was set so that the center of the open field measured 300 +/- 5 lux. A calibration video was taken using a poster board on the bottom of the arena with the center and each corner (1 inch from both edges) marked by circles drawn with black marker. Subjects were initially placed in the center of the arena facing away from the movable cart. The video recording software was started and the subject was allowed to freely explore the arena. After 30 minutes of uninterrupted exploration, the subject was returned to its home cage and the number of fecal boli in the arena was counted. The open field was cleaned with 70% ethanol and five minutes were allotted for the ethanol to evaporate before testing the cagemate.

### Light/Dark Test

A mouse-sized place-preference chamber (27” W x 8” D x 15” H, San Diego Instruments) was used for a light/dark test. One half of the arena had a black floor and was exteriorly covered with black static cling film. The opposite chamber had a white floor and was interiorly covered with white static cling film (except for the lid to allow video observation). The chamber divider was positioned 4 cm above the floor to allow free access between both chambers. Lighting was set so that the center of the light chamber measured 300 +/- 5 lux. A calibration video was taken using a separate floor panel placed on top of the floor in the white chamber with the center and each corner (0.5 inch from both edges) marked by circles drawn with black marker. Subjects were initially placed in the center of the light chamber facing the entrance to the dark chamber. The video recording software was started and the subject was allowed to freely explore the arena. After 30 minutes of uninterrupted exploration, the subject was returned to its home cage and the number of fecal boli in each chamber was counted. Both chambers were cleaned with 70% ethanol and the chamber floors were removed and shaken to evaporate the ethanol before testing the cagemate.

### Predator Cue Test

A rat-sized y-maze (San Diego Instruments) was used for a predator cue test. Walls of the y-maze were increased to 20 inches to prevent escape during testing. Lighting was set so that the center of the y-maze measured 300 +/- 5 lux. Two arms were designated as cue arms where either a predator or control cue would be placed. The third arm was designated as the “blank arm”. Due to the lighting arrangement of the room, the blank arm was darker (∼150 lux) than the center or cue-containing arms. A calibration video was taken using tightly balled black garbage bags inside the ports of each cue arm and a poster board in the blank arm marking the center and edge of the blank arm with black marker. Subjects were initially placed in the center of the empty y-maze facing the blank arm. The video recording software was started and the subject was allowed to freely explore the arena. After 10 minutes, the subject was corralled to the end of the blank arm and a barrier was inserted isolating the subject. Fecal boli were then counted and removed. A cotton ball soaked in 4 ml of red fox urine (Minnesota Traplines) was placed in the port of a randomly chosen cue arm as a predator-associated cue. A cotton ball soaked in 4 ml of white doe urine (Code Blue) was placed in the port of the other cue arm as a non-predator-associated cue. The barrier was then removed and the video recording software was started again. After 20 minutes, the subject was corralled to the end of the blank arm and contained using an insertable barrier. The subject was then anaesthetized using a cassette stuffed with isoflurane-soaked gauze and transferred to a glass jar along with the cassette for blood collection. The cotton balls were removed from the y-maze and fecal boli were counted. The y-maze was cleaned with 70% ethanol and five minutes were allotted for the ethanol to evaporate. The cagemate was then tested using the same procedure and cotton balls.

### Analysis of Videos

Videos were translated into x and y coordinates of the subject for each frame of the video using the following pipeline. First, the raw videos were converted using ffmpeg via the following command:

ffmpeg -i inputFileName.avi -pix_fmt nv12 -f avi -vcodec rawvideo convertedFileName.avi

The converted video file was run through an ImageJ script associated with Mousemove [29], which we modified to label frames deleted due to inability to maintain the framerate or frames when no mouse was detected. Deleted frames and frames when multiple objects were detected were reincorporated using the position of flanking frames. Calibration videos were run through the same process to obtain positions of known locations in the arena. A small subset of videos had a large number of frames (>1%) deleted by erroneously tracking multiple objects. The trajectory files for these videos were manually edited to remove objects that never moved (i.e. not the mouse).

Combined trajectory files were then analyzed to measure a variety of different traits depending on the test (see below). Pixels were translated to mm using known distances between objects in the calibration trajectory file. A mobile threshold was set for each subject using the method of Shoji [30]. Output for each trait was recorded for every minute of the test to observe temporal patterns during the test.

The center of the open field was defined as a circle with diameter half the length of a side (29cm). Distance traveled was the sum of all positional changes above the mobile threshold in the combined trajectory file.

Times spent past three different thresholds in the light chamber were recorded: 1) 0.5 inches from the dark chamber, 2) midpoint of the light chamber, and 3) 0.5 inches from the edge of the wall of the light chamber.

For the predator cue test, entrance into each arm was recorded once a mouse was 12 inches from the end of the arm. Distance traveled was computed as the sum of all positional changes above the mobile threshold in the combined trajectory file.

### Corticosterone Quantification

Immediately following the predator cue test, each mouse was anaesthetized using an isoflurane-soaked gauze and transferred to a glass jar. Once unresponsive, the mouse was decapitated, and blood was collected and stored on ice until all tests for the day were completed. Blood was centrifuged at 1200 rpm for 10 minutes at 4°C to collect plasma. Samples were stored at -80°C until assay submission. Plasma corticosterone concentration was quantified by ELISA at the Wisconsin National Primate Research Center.

### Statistical Analyses

All statistical analyses were conducted in R (v 3.4.1) [31]. Linear models were built separately for each behavior using the lm function {stats}. Behaviors were treated as dependent variables. Strain, sex, age (for the open field test), and cross direction (for F1s) were treated as fixed, independent variables. Interactions suggested by graphical patterns were also evaluated. The significance of an independent variable was evaluated using additional sum-of-squares tests comparing models that included or excluded the variable. Behaviors in F1s and parental strains were compared using t-tests. F1s from mothers of different strains were tested separately when the cross direction term in the linear model was significant. F1s were inferred to be similar to a parental strain when they were statistically indistinguishable from that strain but distinct from the other parental strain. F1s were inferred to be similar to the mid-parent value when they were indistinguishable from that value but distinct from both parental strains. When F1s were statistically distinct from both parental distributions and the mid-parent value they were inferred to be similar to midpoint values between the mid-parent value and the parental mean to which they were closest.

### Data Availability

All computational scripts used in video analysis are available in Supplementary Files 1-9. Raw measurements and associated metadata for each mouse are included in Supplementary Table 1.

## Results

### Open Field Test

We conducted open field tests at two life stages (juvenile and adult) to investigate how GI mice and mainland mice explore a brightly lit, open environment with high risk of detection. We observed extensive behavioral differences between GI mice and mainland mice at both juvenile and adult ages (Figure 2; Table 1). Overall, GI mice travel more, spend more time in the center of the arena, and deposit fewer fecal boli than mainland mice (Figure 2; Table 1). Strain differences for time spent in the center of the arena and the number of fecal boli deposited are similar between ages. Alternatively, distance traveled shows a strong strain-by-age interaction, with the disparity between strains expanding after puberty.

**Table 1:** Summaries of linear model results for each trait across experiments. The third column shows transformations performed on the dependent variable. The fourth column shows significant independent variables among Sex, Strain, and Age (for traits from the open field test). When none of these predictors were significant, the overall mean is presented as the Intercept Estimate.

| Experiment | Trait | Transformation | Independent Variables | Intercept Estimate | Strain (Mainland) Effect |
| --- | --- | --- | --- | --- | --- |
| Open Field | Distance Traveled (meters) | None | Sex, Strain, Age, Strain*Age | 201.686 | -17.416 |
|  | Time Spent Mobile (seconds) | None | Sex, Strain, Age, Strain*Age | 648.324 | -6.506 |
|  | Time in Center (seconds) | Natural Log | Strain | 5.16728 | -0.91655 |
|  | Number of Fecal Boli | None | Strain | 6.833 | 2.7137 |
| Light/Dark Box | Time in Light Chamber (seconds) | None | Strain | 774.53 | -271.62 |
|  | Light Chamber Entrances | Natural Log | Strain | 5.52242 | -0.678 |
|  | Time Past Center (seconds) | None | Strain | 409.12 | -155.35 |
|  | Center Crosses | Natural Log | Strain | 4.63887 | -1.10467 |
|  | Number of Fecal Boli | None | Strain | 6.9697 | 2.5829 |
| Predator Cue | Ratio of Time in Cue Arm/Blank Arm Pre-exposure | None | None | 0.7235549 | - |
|  | Ratio of Time in Control Arm/Blank Arm Pre-exposure | None | None | 0.7340024 | - |
|  | Distance Traveled (meters) Pre-exposure | None | Strain | 80.37 | -28.213 |
|  | Time Spent Mobile (seconds) Pre-exposure | None | Strain | 186.497 | -40.137 |
|  | Ratio of Time in Cue Arm/Blank Arm Post-exposure | None | None | 0.5019955 | - |
|  | Ratio of Time in Control Arm/Blank Arm Post-exposure | None | None | 0.6987754 | - |
|  | Distance Traveled (meters) Post-exposure | None | Strain, Sex | 131.017 | -35.684 |
|  | Time Spent Mobile (seconds) Post-exposure | Natural Log | Strain, Sex | 5.86979 | -0.14411 |
|  | Corticosterone Concentration (ng/mL) | None | Sex | 715.63 | - |

| Strain p-value | Sex (Male) Effect | Sex p-value | Age (Adult) Effect | Age p-value | Strain:Age Effect | Strain:Age p-value |
| --- | --- | --- | --- | --- | --- | --- |
| 0.0204 | -17.896 | 0.00135 | 62.007 | 1.80E-14 | -116.698 | 2.00E-16 |
| 0.74908 | -44.306 | 0.00351 | 99.081 | 2.82E-06 | -245.43 | 6.66E-15 |
| 2.00E-16 | - | - | - | - | - | - |
| 1.79E-14 | - | - | - | - | - | - |
| 6.25E-10 | - | - | - | - | - | - |
| 9.76E-10 | - | - | - | - | - | - |
| 2.54E-07 | - | - | - | - | - | - |
| 2.00E-16 | - | - | - | - | - | - |
| 9.97E-05 | - | - | - | - | - | - |
| - | - | - | - | - | - | - |
| - | - | - | - | - | - | - |
| 1.21E-13 | - | - | - | - | - | - |
| 2.57E-09 | - | - | - | - | - | - |
| - | - | - | - | - | - | - |
| - | - | - | - | - | - | - |
| 4.91E-08 | -17.732 | 0.00374 | - | - | - | - |
| 0.000247 | -0.10018 | 0.010094 | - | - | - | - |
| - | -100.01 | 0.0481 | - | - | - | - |

**Figure 2:**
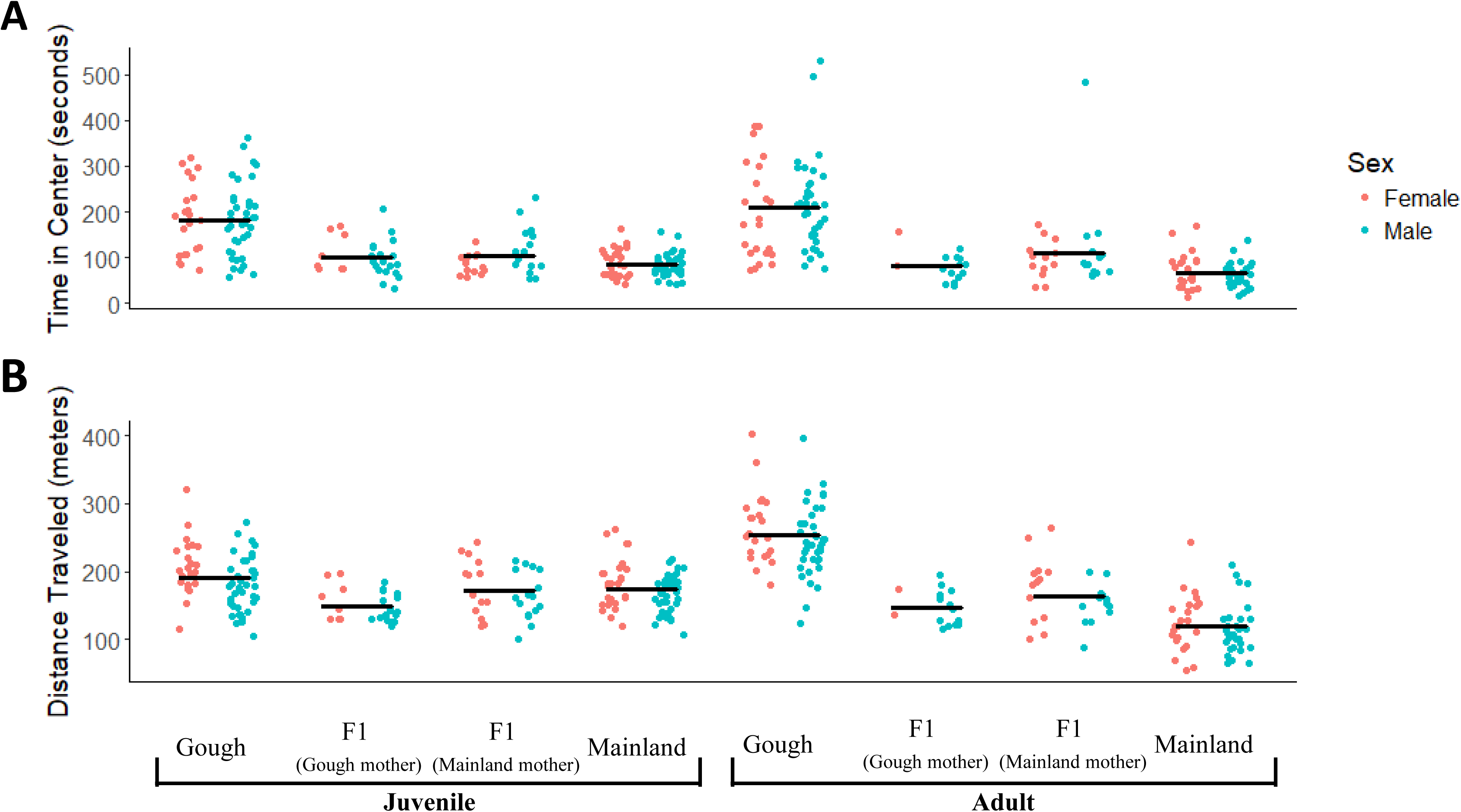
Open field comparisons between strain and ages. Means across sexes are designated by horizontal black bars. F1s are separated based on strain of the mother. A) Time spent in the center of the open field. B) Distance traveled in meters.

F1 offspring from crosses between GI mice and mainland mice show behavioral patterns consistent with a nonadditive genetic architecture with occasional parental effects (Figure 2; Table 2). For most behaviors, F1 averages are closer to the mainland strain. Only the number of fecal boli deposited by F1 adults is statistically indistinguishable from the mid-parent value (Table 2). Cross direction influences distance traveled in both juveniles and adults. F1s with a GI mother travel less than F1s with a mainland mother (Supplementary Table 2).

**Table 2:** Summary of effects influencing behaviors in the open field. A “+” indicates significance of that variable for the given trait. F1 groupings are based on which values (mainland mean, GI mean, or midparent) the F1 distribution is significantly different from using a t-test. For traits with a significant Cross Direction effect the two groups were treated separately.

| Experiment | Trait | Sex Effect | Cross Direction Effect | F1 Grouping |
| --- | --- | --- | --- | --- |
| Open Field Juvenile | Distance Traveled | + | + | Underdominant (GI Mother), Mainland (Mainland Mother) |
|  | Time in Center | - | - | Midparent/Mainland midpoint |
|  | Number of Fecal Boli | - | - | Gough Island |
| Open Field Adult | Distance Traveled | + | + | Midparent/Mainland midpoint |
|  | Time in Center | - | - | Midparent/Mainland midpoint |
|  | Number of Fecal Boli | - | - | Midparent |

Collectively, these results indicate that GI mice evolved an increased willingness to explore novel, risky environments. Whereas boldness-related behaviors are consistent across life stages and cross directions, exploratory behaviors are influenced by age and parental effects.

### Light/Dark Test

To understand how exploration of a novel environment changes when a sheltered area is available, we conducted a light/dark test using a place preference chamber with equally sized light and dark chambers. We found that GI mice spend nearly twice as much time in the light chamber as mainland mice (Figure 3; Table 1). GI mice also enter the light chamber nearly twice as many times. Strain differences are maintained when considering only exploratory bouts past the center of the light chamber. As with the open field test, GI mice deposited less fecal boli, suggesting that baseline anxiety is similar in the two tests (Table 1). These patterns indicate that GI mice evolved increased willingness to leave sheltered areas and to venture farther from shelters compared to mainland mice.

**Figure 3:**
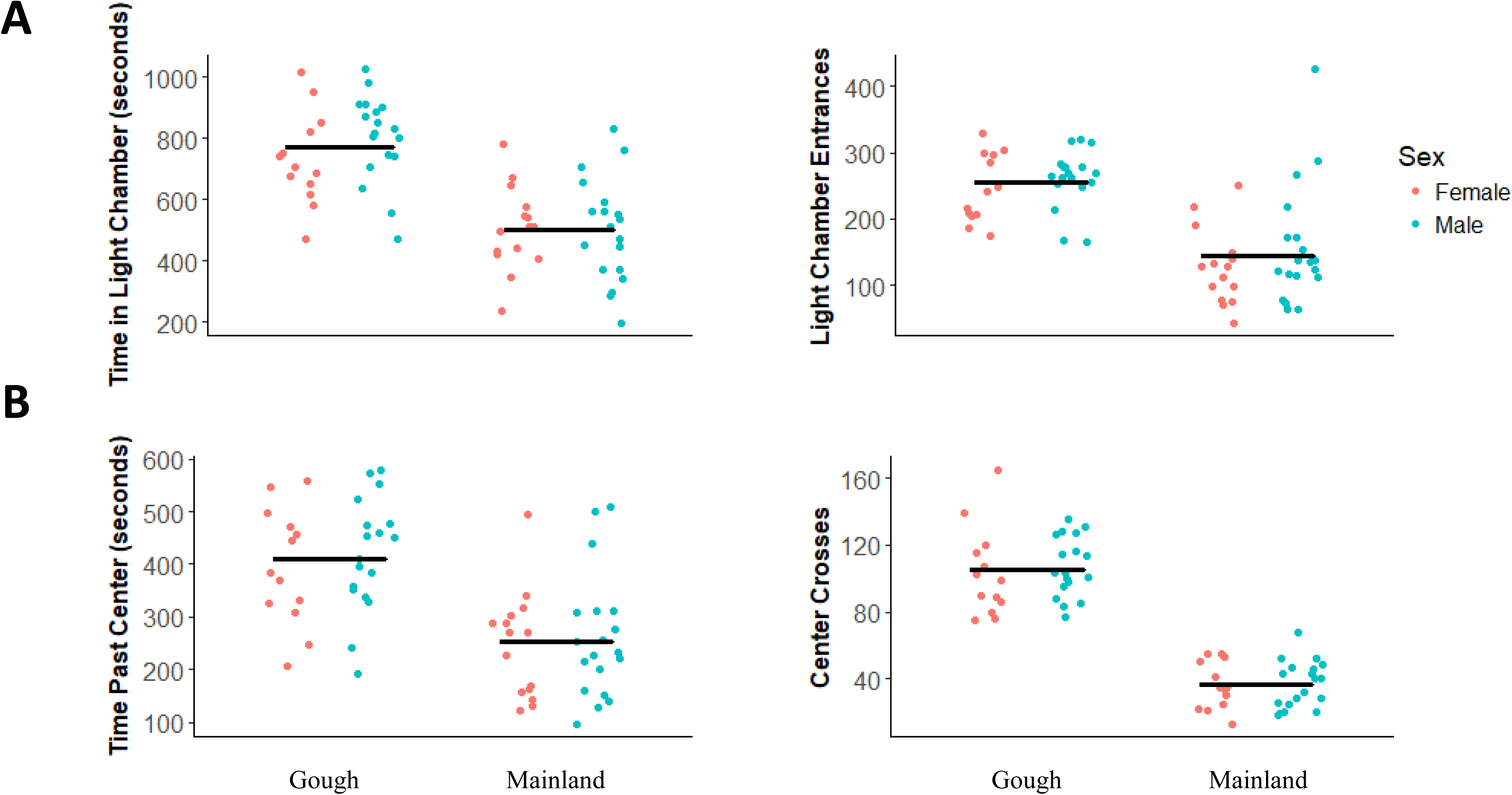
Results from light/dark test. Means across sexes are designated by horizontal black bars. A) Left: amount of time spent in light chamber of light/dark box. Right: number of entrances to the light chamber. B) Left: amount of time spent past the center of the light chamber. Right: number of crosses past the center of the light chamber.

### Predator Cue Test

To reveal how exploration is impacted by the presence of a predator cue, we conducted tests in a y-maze with one arm containing a cotton ball soaked in fox urine. In the first 10 minutes of the test with no cues present, both strains show a slight preference for the blank arm where no cue would be placed (Figure 4A). This pattern could reflect the slight shading of this arm due to lighting constraints in the room. GI mice travel farther overall and deposit fewer fecal boli than mainland mice during these first ten minutes, echoing results from the open field test (Table 1). After both cues are presented, GI mice and mainland mice spend less time in the arm containing the fox urine than in the arm containing the deer urine (Figure 4A; Table 1). We find no evidence for strain differences in this avoidance of the predator cue.

**Figure 4:**
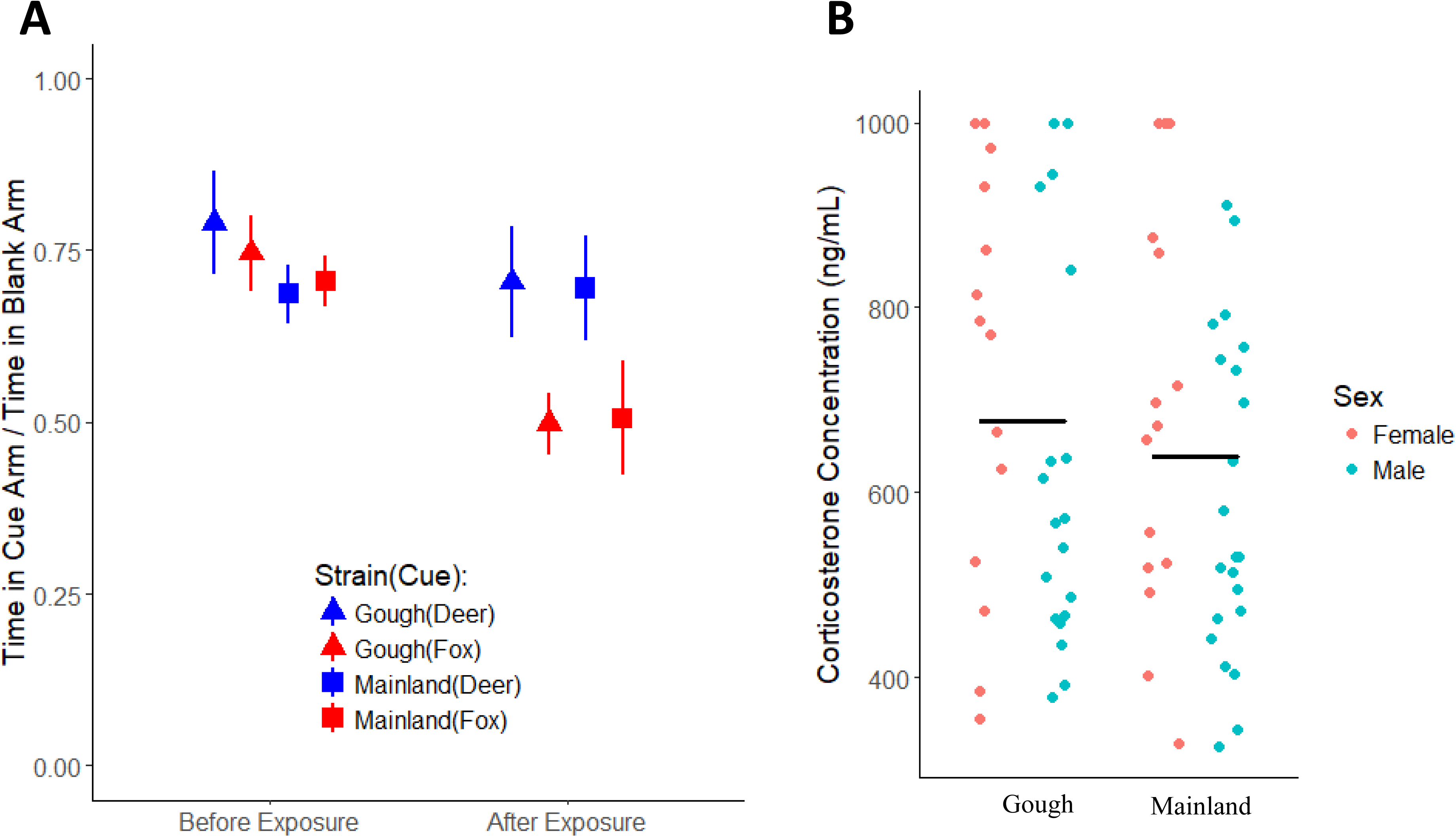
Results from Predator Cue Test. A) Mean ratio and standard error of time spent in arm with a given cue to time spent in blank arm. Ratios on the left are for the ten minutes of exploration before cues were added. B) Corticosterone concentration of mice immediately after the predator cue test. Means across sexes are designated by horizontal bars.

Plasma corticosterone concentrations collected immediately following the predator cue test do not differ between strains (Figure 4B, *t*-test; *P*-value = 0.447). This result indicates that GI mice are as physiologically stressed as mainland mice by the presence of a predator cue though they continue to deposit fewer fecal boli (*t*-test; *P*-value < 1.2E-7). Despite the apparent similarity in anxiety among the two strains, GI mice travel more than mainland mice after cues are presented (Table 1). These findings suggest that GI mice retain avoidance and stress responses associated with exposure to threats from terrestrial predators even while evolving a greater willingness to explore.

## Discussion

Gough Island mice are more exploratory and less fearful than their mainland counterparts in novel, open environments without cues from terrestrial predators. Data from across our experiments demonstrate that GI mice show constant motion throughout each test with little preference for location except when direct cues of terrestrial predators are present. In contrast, mainland mice have high innate anxiety and prefer to remain in regions of each arena where a predator would have the smallest chance of detecting and/or grasping them (e.g. periphery of the open field, dark chamber of the light/dark box). Since all mice in this study were raised in a common setting, these behavioral differences among inbred strains have a genetic basis.

Although understanding which aspects of the environment on Gough Island stimulated behavioral evolution will require ecological studies, characteristics of GI mice and the island suggest potential causes. GI mice are the largest wild house mice in the world [23], offering an extreme case of the gigantism commonly observed among island rodents [15-16], Larger bodies demand greater energetic requirements, particularly during winter [32]. The willingness of GI mice to explore could have been driven by expanded caloric demand, especially with a diet that is highly varied and likely opportunistic [22,33]. Additionally, the evolution of boldness may have facilitated the transition to eating seabird chicks, which are a rich source of nutrients during winter when food is scarce and mortality is high [33]. In contrast to mice inhabiting typical mainland environments, less anxious mice on Gough Island can forage in open areas without danger from predators. For these reasons, exploration and boldness likely provide significant advantages to mice on Gough Island.

Comparisons across juvenile and adult life stages suggest that enhanced exploration and reduced anxiety have evolved along distinct developmental trajectories in GI mice. Although both juvenile and adult GI mice spend more time in the center of an open field and deposit fewer fecal boli, GI mice travel farther than mainland mice only after puberty. Results from other studies support the idea that exploration and boldness can be uncoupled. Pumpkinseed fish approach a novel food source and a potential threat differently [34]. Wild-caught starlings show greater escape motivation than hand-reared starlings when placed in a new cage, but the two groups of birds respond similarly to novel objects [35]. Organisms living in a complex environment are expected to benefit from context-specific evolution of behavior [34,36]. Exploration, which is tied to the need for resources, including food and shelter, takes on greater importance when mice leave the nest and care of their mother. Alternatively, reducing unnecessary anxiety may increase both juvenile and adult fitness by reducing negative health consequences from continuous stress [9,37]. Therefore, natural selection could uncouple the developmental timing of exploration and anxiety.

Rodents living on islands without terrestrial predators are expected to lose the avoidance response to predator cues [38]. It is interesting, therefore, that GI mice and mainland mice respond similarly to fox urine. There are several explanations for this finding, none of which are mutually exclusive. First, GI mice may not have been on the island long enough for new genetic variants that relax the response to predator cues to arise and spread. Perhaps this phenotype has a smaller mutational target size than general anxiety, a trait for which we observed a heritable reduction in GI mice. Secondly, the source population(s) of GI mice may have experienced frequent terrestrial predation but not avian predation, leaving intact the response to cues from terrestrial predators while allowing the loss of anxiety about aerial threats. This idea is consistent with the “predator archetype” hypothesis [39], which postulates disparate defense responses to avian and terrestrial predators. A third possibility is that selection on general anxiety may be more direct than selection to remove predator cue responses. The presence or absence of a predator cue response makes no difference when predators are absent, whereas reducing general anxiety can facilitate the optimization of foraging strategies and reduce energy expenditure from unnecessary stress [9]. A final explanation is that the pathway for detecting terrestrial predators could play additional functional roles in GI mice. Sulfur-containing byproducts of meat-eating vertebrates are known to be aversive stimuli for rodents [40], but there may be other sulfur-containing compounds on Gough Island that mice need to detect. Since fish-eating seabirds are a major source of nutrition for GI mice during the winter season [26,33], the ability to locate nests emitting sulfur signals could be beneficial.

The phenotypic patterns we documented in F1s provide insights into the genetic architecture that underlies the evolution of new behaviors in GI mice. We found that F1s resemble mainland mice more than GI mice in open field tests, implying that GI mouse alleles are recessive to mainland mouse alleles for several behaviors connected to boldness and exploration. Recessive mutations tend to reduce function, suggesting that the variants of interest could have decreased the expression of genes or pathways that are typically active in mainland mice. The evolutionary trajectory of recessive mutations in GI mice would have depended on their initial frequencies. Whereas new recessive variants would have required stronger selection to become established and spread to high frequency than new dominant variants [41], the probability of fixation of standing variants that contribute to adaptation is independent of dominance [42]. The inference of recessive gene action also raises the prospect that behavioral evolution was accomplished through a small number of mutations with large phenotypic effects. Although loci that affect behavior in laboratory strains of mice have been mapped to every chromosome [43], alleles with substantial effects exist for some behaviors, including boldness and exploration [44-45].

Our characterization of F1s uncovered additional factors that shape behavior in GI mice. An apparent maternal effect on distance traveled in the open field acts in the opposite direction of the strain effect (i.e. GI mouse mothers reduce distance traveled by F1 offspring). This result implies that the genetic increase in distance traveled in GI mice is greater than it appears based on comparisons to mainland mice since this maternal effect must be overcome. Antagonism between maternal and direct genetic effects provides a barrier to selection and could draw populations away from optimal trait values, depending on the degree of covariance between these effects [46].

Several caveats accompany our interpretations. The mainland strain we profiled was chosen based on its availability and common usage in mouse genetics. If the behavior of this wild-derived inbred strain departs significantly from the mainland mice from which GI mice are descended, our conclusions about the evolution of new behaviors in GI mice could be incorrect. Western Europe is the most likely ancestral origin of *M. m. domesticus* in North America and on Gough Island [20]. Detailed reconstruction of behavioral evolution in GI mice will ultimately require behavioral studies in specific source populations, which remain to be identified. Our study also assumes behavior in the laboratory is representative of behavior in the wild. While the simplicity of our experimental design facilitates the identification of causal factors in behavioral differences, those interpretations are limited to the environment in which the tests were conducted. Mice in the wild experience complex environments without ad libitum access to food. Incorporating more realistic scenarios (e.g. fasting, outside enclosures) in future behavioral studies will help broaden our interpretations of the behavioral changes in GI mice.

Our study is among the first to report inherited differences in behavior between island and mainland populations [17,47]. Island deer mice that evolved larger bodies are less aggressive than their mainland relatives, but this behavioral difference is not heritable [13]. Perhaps the extreme ecological conditions on Gough Island (e.g. lack of predators, lack of human commensals, presence of seabird chicks as a source of food in winter) have created particularly strong selective pressure for behavioral evolution. Regardless of the environmental causes, our demonstration of heritable differences between island and mainland populations in a genetic model organism sets the stage for identifying genes responsible for behavioral evolution associated with island colonization. GI mice could also serve as a useful experimental model for exploration and anxiety in humans [48-49] where the list of candidate genes for these behaviors continues to grow [50].

## Authors’ Contributions

JAS helped design the study, carried out the behavioral testing, performed the video analysis, conducted the statistical testing, and drafted the manuscript; MJN conceived of the study, helped design the study, and critically revised the manuscript; BAP helped design the study, and critically revised the manuscript. All authors gave final approval for publication and agree to be held accountable for the work performed therein.

### Acknowledgments

We thank Megan Latsch and Dr. Michelle Parmenter for conducting the pilot study that inspired this project. We thank Dr. Andrew Bakken for helping construct the open field and Dr. John Stratton for assistance in developing scripts.

## Ethics

All experiments were approved by the University of Wisconsin-Madison Institutional Animal Care and Use Committee (protocol no. V005209).

