## Supplementary File 1 for "Evolution of Boldness and Exploratory Behavior in Giant Mice from Gough Island"

__author__ = 'Jered Stratton'
# Combines trajectory files from Preprocessing.ijm and incorporates deleted frames using average of flanking frame
# coordinates
import sys
import re

trajFileName = sys.argv[1]  # First trajectory file to input
deletedFileName = sys.argv[2]  # File of deleted frame numbers
outFilePrefix = sys.argv[3]  # Prefix for output file
frameTotal = int(sys.argv[4])  # Total number of frames in video

frameStep = 5000  # Number of frames per file
trajFrameCount = 1  # Current frame number for output
frameIndex = 1  # Current frame number for loading files
nullFrames = []  # Frames with more or less than one object detected
nullCur = 0  # Index of last null frame detected
nullBlockSize = 0
trajFrames = []  # Data for combined traj file
outFrames = []  # Data for output traj file with deleted frames

# Obtain prefix for traj file names
trajFilePrefix = re.search('.*(?=\d+-\d+)', trajFileName)

# Function for removing empty tracks
def filterLineValues(lineValue):
    if lineValue == ' ':
        return False
    else:
        return True


# Combine trajectory files from Preprocessing.ijm output
while frameIndex < frameTotal:
    fend = min(frameIndex + frameStep - 1, frameTotal)
    trajFileName = trajFilePrefix.group(0) + str(frameIndex) + '-' + str(fend) + '.txt'
    trajFile = open(trajFileName, 'r')
    trajFile.readline()
    trajFile.readline()
    for line in trajFile:
        line = line.rstrip()
        lineValues = line.split('\t')
        lineValues = list(filter(filterLineValues, lineValues))
        if len(lineValues) > 3:
            if lineValues[3] == '*':
                lineValues.remove('*')
        if len(lineValues) > 3:	
            print("More than one object detected at frame " + str(trajFrameCount))
            trajFrames.append([trajFrameCount, None, None])
        elif len(lineValues) > 1:
            trajFrames.append([trajFrameCount, float(lineValues[1]), float(lineValues[2])])
        else:
            print("Null frame detected at frame " + str(trajFrameCount))
            trajFrames.append([trajFrameCount, None, None])
        trajFrameCount += 1
    frameIndex += frameStep
    trajFile.close()

# Record deleted frames
print(str(len(trajFrames))+" frames found in combined TkResults files")
outIndex = 1
trajIndex = 0
deletedFile = open(deletedFileName, 'r')
deletedFile.readline()
blockIndex = re.search('\d+(?=\.\d+)', deletedFile.readline())  # Frame number where block is incorporated
while int(blockIndex.group(0)) == 0:
    blockIndex = re.search('\d+(?=\.\d+)', deletedFile.readline())
deletedBlockSize = 1  # Number of frames in block
for line in deletedFile:
    deletedCur = re.search('\d+(?=\.\d+)', line)
    # Add line to block of deleted Frames if matches blockIndex
    if deletedCur.group(0) == blockIndex.group(0):
        deletedBlockSize += 1
    # If line is start of new block
    else:
        # Copy trajFrames to outFrames up to deletedBlock
        while int(blockIndex.group(0)) >= trajIndex+1:
            outFrames.append([outIndex, trajFrames[trajIndex][1], trajFrames[trajIndex][2]])
            outIndex += 1
            trajIndex += 1
        for i in range(0, deletedBlockSize):
            outFrames.append([outIndex, None, None])
            outIndex += 1
        blockIndex = deletedCur
        deletedBlockSize = 1

if blockIndex is not None:
    while int(blockIndex.group(0)) >= trajIndex+1:
        outFrames.append([outIndex, trajFrames[trajIndex][1], trajFrames[trajIndex][2]])
        outIndex += 1
        trajIndex += 1
    for i in range(0, deletedBlockSize):
        outFrames.append([outIndex, None, None])
        outIndex += 1
while len(trajFrames) >= trajIndex+1:
    outFrames.append([outIndex, trajFrames[trajIndex][1], trajFrames[trajIndex][2]])
    outIndex += 1
    trajIndex += 1
deletedFile.close()

# Interpolate position of missing frames
# Trim deleted frames at beginning of trajectory file
blockIndex = outFrames[0][1]
while blockIndex is None:
    print("Deleted frame at beginning of video")
    outFrames.pop(0)
    blockIndex = outFrames[0][1]
xPrev = outFrames[0][1]
yPrev = outFrames[0][2]
for i in range(0, len(outFrames)):
    # Add one to nullBlockSize if deleted frame is found
    if outFrames[i][1] is None:
        nullBlockSize += 1
    else:
        # Assign coordinates to deleted block based on flanking frame coordinates
        xNext = outFrames[i][1]
        yNext = outFrames[i][2]
        xStep = (xNext - xPrev) / (nullBlockSize + 1)
        yStep = (yNext - yPrev) / (nullBlockSize + 1)
        for j in range(0, nullBlockSize):
            outFrames[i-nullBlockSize+j] = [i-nullBlockSize+j+1, xPrev+(xStep*(j+1)), yPrev+(yStep*(j+1))]
        nullBlockSize = 0
        xPrev = xNext
        yPrev = yNext
# Trim deleted frames at end of trajectory file
blockIndex = outFrames[len(outFrames)-1][1]
while blockIndex is None:
    print("Deleted frame at end of video")
    outFrames.pop(len(outFrames)-1)
    blockIndex = outFrames[len(outFrames)-1][1]

# Output frames to txt file
outFile = open(outFilePrefix + "_1-" + str(len(outFrames)) + ".txt", "w")
outFile.write("Frame\tX1\tY1\tFlag1\n")
outFile.write("Tracks 1 to 1\n")
for i in range(0, len(outFrames)):
    outFile.write("\t".join(str(x) for x in outFrames[i]))
    outFile.write("\n")
outFile.close()
