## Supplementary File 4 for "Evolution of Boldness and Exploratory Behavior in Giant Mice from Gough Island"

__author__ = 'Jered Stratton'
# Calculates amount of time spent in light chamber of light/dark box
import sys
import math
import statistics
import re

trajFileName = sys.argv[1]  # First trajectory file to input
calibrationFileName = sys.argv[2]
deletedFileName = sys.argv[3]  # File of deleted frame numbers
outFilePrefix = sys.argv[4]  # Prefix for output file
frameTotal = int(sys.argv[5])  # Total number of frames in video

outFileName = outFilePrefix + "_lightDarkData.csv"
output = []
frameStep = 5000  # Number of frames per file
trajFrameCount = 1  # Current frame number for output
frameIndex = 1  # Current frame number for loading files
nullFrames = []  # Frames with more or less than one object detected
nullCur = 0  # Index of last null frame detected
nullBlockSize = 0
trajFrames = []  # Data for combined traj file
outFrames = []  # Data for output traj file with deleted frames
xPrev = 0
yPrev = 0
frameCount = 0
timeFactor = 1/30  # Number of seconds per frame
reportInterval = 60/timeFactor  # Number of frames before reporting measurements again
timePastRight = 0
timePastCenter = 0
timePastLeft = 0
rightEntrances = 0
centerEntrances = 0
leftEntrances = 0
rightLatency = 0
centerLatency = 0
leftLatency = 0
pastRight = False
pastCenter = False
pastLeft = False

# Shorthand for positions in calibration file
centerX = 0
centerY = 0
BRCx = 0
BRCy = 0
BLCx = 0
BLCy = 0
TRCx = 0
TRCy = 0
TLCx = 0
TLCy = 0

# Read in the calibration file and store positions in dummy objects
dummyX = []
dummyY = []
for i in range(1, 6):
    xPositions = []
    yPositions = []
    calibrationFile = open(calibrationFileName, 'r')
    calibrationFile.readline()
    calibrationFile.readline()
    for line in calibrationFile:
        line = line.rstrip()
        lineValues = line.split('\t')
        xPositions.append(float(lineValues[(i*3)-2]))
        yPositions.append(float(lineValues[(i*3)-1]))
    dummyX.append(statistics.mean(xPositions))
    dummyY.append(statistics.mean(yPositions))
    calibrationFile.close()

# Identify center
centerX = statistics.median(dummyX)
centerY = statistics.median(dummyY)
dummyX.remove(statistics.median(dummyX))
dummyY.remove(statistics.median(dummyY))

# Identify corners based on position relative to center
for i in range(0, 4):
    xCur = dummyX[i]
    yCur = dummyY[i]
    if xCur > centerX and yCur > centerY:
        BRCx = xCur
        BRCy = yCur
    elif xCur < centerX and yCur > centerY:
        BLCx = xCur
        BLCy = yCur
    elif xCur > centerX and yCur < centerY:
        TRCx = xCur
        TRCy = yCur
    elif xCur < centerX and yCur < centerY:
        TLCx = xCur
        TLCy = yCur

rightDistance = math.sqrt((TRCx - BRCx) ** 2 + (TRCy - BRCy) ** 2)  # Distance between two right points
leftDistance = math.sqrt((TLCx - BLCx) ** 2 + (TLCy - BLCy) ** 2)  # Distance between two left points
centerRightDistance = abs((BRCy-TRCy)*centerX-(BRCx-TRCx)*centerY+(BRCx*TRCy)-(BRCy*TRCx))/rightDistance  # Distance from center to right threshold
centerLeftDistance = abs((BLCy-TLCy)*centerX-(BLCx-TLCx)*centerY+(BLCx*TLCy)-(BLCy*TLCx))/leftDistance  # Distance from center to left threshold
leftRightDistance = centerRightDistance + centerLeftDistance

# Obtain prefix for traj file names
trajFilePrefix = re.search('.*(?=\d+-\d+)', trajFileName)

# Function for removing empty tracks
def filterLineValues(lineValue):
    if lineValue == ' ':
        return False
    else:
        return True

# Function for collecting data in each frame
def collectFrameInfo(xCur, yCur):
    global pastRight
    global pastCenter
    global pastLeft
    global rightEntrances
    global centerEntrances
    global leftEntrances
    global rightLatency
    global centerLatency
    global leftLatency
    global timePastRight
    global timePastCenter
    global timePastLeft
    global output
    global frameCount
    global frameTotal
    mRight = abs((BRCy - TRCy) * xCur - (BRCx - TRCx) * yCur + (BRCx * TRCy) - (BRCy * TRCx)) / rightDistance
    mLeft = abs((BLCy - TLCy) * xCur - (BLCx - TLCx) * yCur + (BLCx * TLCy) - (BLCy * TLCx)) / leftDistance
    if mLeft > leftRightDistance:  # Mouse is on right of right threshold
        pastRight = False
        pastCenter = False
        pastLeft = False
    else:
        if mRight <= mLeft:  # Mouse is between right and center thresholds
            if not pastRight:
                rightEntrances += 1
            pastRight = True
            if rightLatency == 0:
                rightLatency = frameCount
                print(str(rightLatency * timeFactor) + " seconds to cross right threshold for the first time")
            pastCenter = False
            pastLeft = False
            timePastRight += 1
        else:  # Mouse is left of center threshold
            if mRight > leftRightDistance:  # Mouse is past left threshold
                if not pastLeft:
                    leftEntrances += 1
                pastLeft = True
                pastCenter = True
                pastRight = True
                if leftLatency == 0:
                    leftLatency = frameCount
                    print(str(leftLatency * timeFactor) + " seconds to cross left threshold for the first time")
                    if centerLatency == 0:
                        centerLatency = frameCount
                        print(str(centerLatency * timeFactor) + " seconds to cross center threshold for the first time")
                        rightLatency = frameCount
                        print(str(rightLatency * timeFactor) + " seconds to cross right threshold for the first time")
                timePastRight += 1
                timePastCenter += 1
                timePastLeft += 1
            else:  # Mouse is between center and left thresholds
                if not pastCenter:
                    centerEntrances += 1
                pastCenter = True
                pastRight = True
                pastLeft = False
                if centerLatency == 0:
                    centerLatency = frameCount
                    print(str(centerLatency * timeFactor) + " seconds to cross center threshold for the first time")
                    if rightLatency == 0:
                        rightLatency = frameCount
                        print(str(rightLatency * timeFactor) + " seconds to cross right threshold for the first time")
                timePastRight += 1
                timePastCenter += 1
    if frameCount % reportInterval == 0:
        output.append([(frameCount*timeFactor), (timePastRight*timeFactor), (timePastCenter*timeFactor), (timePastLeft*timeFactor), rightEntrances, centerEntrances, leftEntrances])


# Combine trajectory files from Preprocessing.ijm output
while frameIndex < frameTotal:
    fend = min(frameIndex+frameStep-1, frameTotal)
    trajFileName = trajFilePrefix.group(0) + str(frameIndex) + '-' + str(fend) + '.txt'
    trajFile = open(trajFileName, 'r')
    trajFile.readline()
    trajFile.readline()
    for line in trajFile:
        line = line.rstrip()
        lineValues = line.split('\t')
        lineValues = list(filter(filterLineValues, lineValues))
        if len(lineValues) > 3:
            if lineValues[3] == '*':
                lineValues.remove('*')
        if len(lineValues) > 3:	
            print("More than one object detected at frame " + str(trajFrameCount))
            trajFrames.append([trajFrameCount, None, None])
        elif len(lineValues) > 1:
            trajFrames.append([trajFrameCount, float(lineValues[1]), float(lineValues[2])])
        else:
            print("Null frame detected at frame " + str(trajFrameCount))
            trajFrames.append([trajFrameCount, None, None])
        trajFrameCount += 1
    frameIndex += frameStep
    trajFile.close()

# Record deleted frames
outIndex = 1
trajIndex = 0
deletedFile = open(deletedFileName, 'r')
deletedFile.readline()
blockIndex = re.search('\d+(?=\.\d+)', deletedFile.readline())  # Frame number where block is incorporated
while int(blockIndex.group(0)) == 0:
    outFrames.append([outIndex, None, None])
    blockIndex = re.search('\d+(?=\.\d+)', deletedFile.readline())
deletedBlockSize = 1  # Number of frames in block
for line in deletedFile:
    deletedCur = re.search('\d+(?=\.\d+)', line)
    # Add line to block of deleted Frames if matches blockIndex
    if deletedCur.group(0) == blockIndex.group(0):
        deletedBlockSize += 1
    # If line is start of new block
    else:
        # Copy trajFrames to outFrames up to deletedBlock
        while int(blockIndex.group(0)) >= trajIndex+1:
            outFrames.append([outIndex, trajFrames[trajIndex][1], trajFrames[trajIndex][2]])
            outIndex += 1
            trajIndex += 1
        for i in range(0, deletedBlockSize):
            outFrames.append([outIndex, None, None])
            outIndex += 1
        blockIndex = deletedCur
        deletedBlockSize = 1

if blockIndex is not None:
    while int(blockIndex.group(0)) >= trajIndex+1:
        outFrames.append([outIndex, trajFrames[trajIndex][1], trajFrames[trajIndex][2]])
        outIndex += 1
        trajIndex += 1
    for i in range(0, deletedBlockSize):
        outFrames.append([outIndex, None, None])
        outIndex += 1
while len(trajFrames) >= trajIndex+1:
    outFrames.append([outIndex, trajFrames[trajIndex][1], trajFrames[trajIndex][2]])
    outIndex += 1
    trajIndex += 1
deletedFile.close()

# Collect data from combined list of frames
print("Analyzing video at " + str(1/timeFactor) + " frames per second")
frameCount = 1
nullBlockSize = 0
xPrev = outFrames[0][1]
yPrev = outFrames[0][2]
for i in range(0, len(outFrames)):
    xCur = outFrames[i][1]
    yCur = outFrames[i][2]
    # Add 1 to block of deleted Frames if blank frame found
    if xCur is None:
        nullBlockSize += 1
    # If line is not deleted
    else:
        if xPrev is None:  # Deleted Block found at beginning of video
            curLeft = abs((BLCy - TLCy) * xCur - (BLCx - TLCx) * yCur + (BLCx * TLCy) - (BLCy * TLCx)) / leftDistance
            if curLeft <= leftRightDistance and nullBlockSize >= 5:
                print("Warning: deleted block of size " + str(nullBlockSize) + " detected past right threshold at frame: " + str(frameCount))
            for j in range(0, nullBlockSize):
                collectFrameInfo(xCur, yCur)
                frameCount += 1
            xPrev = xCur
            yPrev = yCur
            nullBlockSize = 0
        # Add coordinates to deleted block based on flanking frame coordinates and add to output
        prevLeft = abs((BLCy - TLCy) * xPrev - (BLCx - TLCx) * yPrev + (BLCx * TLCy) - (BLCy * TLCx)) / leftDistance
        curLeft = abs((BLCy - TLCy) * xCur - (BLCx - TLCx) * yCur + (BLCx * TLCy) - (BLCy * TLCx)) / leftDistance
        if (prevLeft <= leftRightDistance and curLeft <= leftRightDistance) and nullBlockSize >= 5:
            print("Warning: deleted block of size " + str(nullBlockSize) + " detected past right threshold at frame: " + str(frameCount))
        if curLeft > prevLeft:
            xPrev = xCur
            yPrev = yCur
        for j in range(0, nullBlockSize):
            collectFrameInfo(xPrev, yPrev)
            frameCount += 1
        collectFrameInfo(xCur, yCur)
        frameCount += 1
        xPrev = xCur
        yPrev = yCur
        nullBlockSize = 0

if xCur is None:
    prevLeft = abs((BLCy - TLCy) * xPrev - (BLCx - TLCx) * yPrev + (BLCx * TLCy) - (BLCy * TLCx)) / leftDistance
    if prevLeft <= leftRightDistance and nullBlockSize >= 5:
        print("Warning: deleted block of size " + str(nullBlockSize) + " detected past right threshold at frame: " + str(frameCount))
    for j in range(0, nullBlockSize):
        collectFrameInfo(xPrev, yPrev)
        frameCount += 1

output.append([(frameCount*timeFactor), (timePastRight*timeFactor), (timePastCenter*timeFactor), (timePastLeft*timeFactor), rightEntrances, centerEntrances, leftEntrances])

# Output data to csv file
outFile = open(outFileName, "w")
outFile.write("Time (seconds),Time Past Right Threshold (seconds), Time Past Center Threshold (seconds), Time Past Left Threshold (seconds), Number of Right Threshold Crosses, Number of Center Threshold Crosses, Number of Left Threshold Crosses\n")
for i in range(0, len(output)):
    outFile.write(",".join(str(x) for x in output[i]))
    outFile.write("\n")
outFile.close()
