## Supplementary File 3 for "Evolution of Boldness and Exploratory Behavior in Giant Mice from Gough Island"

__author__ = 'Jered Stratton'
# Takes in trajectory file output from Preprocessing.ijm and calculates a variety of behavioral measures reported
# regularly at a specified time interval
# ROI: region of interest
import sys
import math
import statistics
import numpy
import calc_mobile_thresh as cmt


trajFileName = sys.argv[1]  # Combined trajectory file to analyze
calibrationFileName = sys.argv[2]  # Trajectory file of calibration
outFilePrefix = sys.argv[3]  # Prefix for output file
outFileName = outFilePrefix + "_openFieldData.csv"

mobileThresh = 0.0  # Distance (in pixels) needed to say a mouse is moving
mobile = False  # Indicator if subject is mobile
timeFactor = 1/30  # Number of seconds per frame ***CHANGE THIS***
reportInterval = 60/timeFactor  # Number of frames before reporting measurements again
inCenter = False  # Indicator if subject was previously in center (for tracking center entrances)

positions = {'center': [0, 0], 'TRC': [0, 0], 'TLC': [0, 0], 'BRC': [0, 0], 'BLC': [0, 0]}  # List of X,Y coordinates (set by calibration)
distanceFactor = 0  # Number of mm per pixel (set by calibration)

frameCount = 1  # Total number of frames recorded
xPrev = 0
yPrev = 0

distanceTravelled = 0  # Distance travelled
centerEntrances = 0  # Number of times the subject entered the center
timeMobile = 0  # Amount of time (in frames) the subject was mobile
timeROI = {'center': 0, 'TRC': 0, 'TLC': 0, 'BRC': 0, 'BLC': 0}  # Amount of time (in seconds) the subject was in each ROI. TRC = top right corner, TLC = top left corner, BRC = bottom right corner, BLC = bottom left corner
radiusROI = {'center': 0, 'TRC': 0, 'TLC': 0, 'BRC': 0, 'BLC': 0}  # Radius for each ROI
mobileLatency = 0  # Amount of time before the subject moved for the first time
centerLatency = 0  # Amount of time before the subject entered the center for the first time

output = []
# Calibrate the positions of each ROI and the distance in pixels for 1 mm
#
# Read in the calibration file and store positions in dummy objects
dummyX = []
dummyY = []
for i in range(1, 6):
    xPositions = []
    yPositions = []
    calibrationFile = open(calibrationFileName, 'r')
    calibrationFile.readline()
    calibrationFile.readline()
    for line in calibrationFile:
        line = line.rstrip()
        lineValues = line.split('\t')
        xPositions.append(float(lineValues[(i*3)-2]))
        yPositions.append(float(lineValues[(i*3)-1]))
    dummyX.append(statistics.mean(xPositions))
    dummyY.append(statistics.mean(yPositions))
    calibrationFile.close()

# Identify center
positions['center'] = [statistics.median(dummyX), statistics.median(dummyY)]
dummyX.remove(statistics.median(dummyX))
dummyY.remove(statistics.median(dummyY))

# Identify corners based on position relative to center
for i in range(0, 4):
    xCur = dummyX[i]
    yCur = dummyY[i]
    if xCur > positions['center'][0] and yCur > positions['center'][1]:
        positions['BRC'][0] = xCur
        positions['BRC'][1] = yCur
    elif xCur < positions['center'][0] and yCur > positions['center'][1]:
        positions['BLC'][0] = xCur
        positions['BLC'][1] = yCur
    elif xCur > positions['center'][0] and yCur < positions['center'][1]:
        positions['TRC'][0] = xCur
        positions['TRC'][1] = yCur
    elif xCur < positions['center'][0] and yCur < positions['center'][1]:
        positions['TLC'][0] = xCur
        positions['TLC'][1] = yCur

# Set distanceFactor and ROI radii
centerDistances = []
for key in positions:
    distanceToCenter = math.sqrt((positions['center'][0] - positions[key][0])**2 + (positions['center'][1] - positions[key][1])**2)
    centerDistances.append(distanceToCenter)
centerDistances.remove(min(centerDistances))
print("Distances to center:")
print(centerDistances)
distanceFactor = 376.05/statistics.mean(centerDistances)
print("Calibration at " + str(distanceFactor) + "mm per pixel")
for key in radiusROI:
    radiusROI[key] = 145.65/distanceFactor

# Set Mobile threshold using method of Shoji et al.
frameCount = 0
mobileThresh, totalFrames = cmt.mobilityThreshold(trajFileName)

print("Analyzing video at " + str(1/timeFactor) + " frames per second")
# Read in trajectory file
trajFile = open(trajFileName, 'r')
trajFile.readline()
trajFile.readline()
for line in trajFile:
    line = line.rstrip()
    lineValues = line.split('\t')
    xCur = float(lineValues[1])
    yCur = float(lineValues[2])
    if frameCount == 0:
        xPrev = xCur
        yPrev = yCur
        # Check for ROI
        for key in positions:
            if math.sqrt((positions[key][0] - xCur)**2 + (positions[key][1] - yCur)**2) <= radiusROI[key]:
                    timeROI[key] += timeFactor
                    if key == "center":
                        inCenter = True
        frameCount += 1
        continue
    else:
        distanceCur = math.sqrt((xCur - xPrev)**2 + (yCur - yPrev)**2)
        if distanceCur > mobileThresh:
            mobile = True
            if mobileLatency == 0:
                mobileLatency = frameCount
                print(str(mobileLatency*timeFactor) + " seconds to move for the first time")
        else:
            mobile = False
        for key in positions:
            if math.sqrt((positions[key][0] - xCur)**2 + (positions[key][1] - yCur)**2) <= radiusROI[key]:
                timeROI[key] += timeFactor
                if key == "center":
                    if not inCenter:
                        centerEntrances += 1
                        if centerLatency == 0:
                            centerLatency = frameCount
                            print(str(centerLatency*timeFactor) + " seconds to enter center")
                    inCenter = True
            elif key == "center":
                inCenter = False
        if mobile:
            distanceTravelled += distanceCur*distanceFactor
            timeMobile += timeFactor
        frameCount += 1
        if frameCount % reportInterval == 0 or frameCount == totalFrames:
            output.append([(frameCount*timeFactor), distanceTravelled/1000, timeMobile, centerEntrances, (timeROI['center']), (timeROI['TLC']), (timeROI['TRC']), (timeROI['BLC']), (timeROI['BRC'])])
        xPrev = xCur
        yPrev = yCur
trajFile.close()

outFile = open(outFileName, "w")
outFile.write("Time (seconds),Distance_Traveled (m),Time_mobile (seconds),Center_Entrances,Time_in_Center,Time_in_TLC,Time_in_TRC,Time_in_BLC,Time_in_BRC\n")
for i in range(0, len(output)):
    outFile.write(",".join(str(x) for x in output[i]))
    outFile.write("\n")
outFile.close()
