## Supplementary File 6 for "Evolution of Boldness and Exploratory Behavior in Giant Mice from Gough Island"

//Preprocessing macro, made by Dr. Lining JU, edited by Jered A. Stratton
run("Close All");
print("\\Clear");
//Indicate the video files locations
HomeDirectory=getDirectory("Choose Source Directory");
list=getFileList(HomeDirectory);
//Reset the background color
run("Colors...", "background=white");
//Setup the stepwise analysis 
framestep=5000;
framerate = 30; // Set this to fps of video
tPrev = 0.03; // Set this to timestamp of first frame based on fps
//Welcome message
Msg="Start batch preprocessing...";
print(Msg);

for (k=0;k<list.length;k++)
{
	ix=indexOf(list[k],"empty");
	if (ix>0)//Check whether this is a background video
	{
		Refname=list[k];
		Expname= substring(Refname,0,ix-1)+".avi";
//Setup the file paths
Refpath=HomeDirectory+File.separator+Refname;
Exppath=HomeDirectory+File.separator+Expname;
Deletedpath=HomeDirectory+File.separator+substring(Expname,0,ix-10)+"Deleted_Frames"+".txt";
Msg="Start processing "+Expname+"...";
print(Msg);
File.append("previous.deleted", Deletedpath);

run("AVI...", "select=[&Exppath] use convert");
framerange=nSlices;
close();

frameCount = 0;
for(i=1;i<=framerange;i=i+framestep)
{
fstart=i;
fend=minOf(i+framestep-1,framerange);
if (fend == (framerange)) {
	fend = 0;
}
m=1; //index of deleted frames - Jered S.
//////////////////////////////////////////////////////////////////
//Import the last image of background avi video
run("AVI...", "select=[&Refpath] use convert");
N=nSlices;
close();
run("AVI...", "select=[&Refpath] first=&N last=&N use convert");

//Import the experiment avi video and select frame numbers
run("AVI...", "select=[&Exppath] first=&fstart last=&fstart use convert");
//Make a stack for alignment
run("Concatenate...", "  title=[Concatenated Alignment] image1=&Expname image2=&Refname image3=[-- None --]");
//Image binarization
setOption("BlackBackground", false);
run("Make Binary", "method=Minimum background=Light thresholded remaining black");
//Align the experiment video with the background video
//run("StackReg ", "transformation=[Rigid Body]");
//Import the experiment avi video and select frame numbers
run("AVI...", "select=[&Exppath] first=&fstart last=&fend use convert");
print(toString(nSlices)+" Frames after import");
//Alignment with the 1st image as the background 
run("Concatenate...", "  title=[Concatenated Stacks] image1=[Concatenated Alignment] image2=[&Expname] image3=[-- None --]");
run("Delete Slice");
//Bin the videos
run("Bin...", "x=2 y=2 z=1 bin=Average");
//Automatically find the boundary edge based on the 1st image.
doWand(160, 115, 90.0, "Legacy");
run("Auto Threshold", "method=Minimum white stack");
//Clear outer area and leave the arena and the mouse.
run("Clear Outside", "stack");

//Delete ROI
run("Select None");
//Remove the first slice
run("Delete Slice");
//Smooth the tracked object
run("Minimum...", "radius=4  stack");

j=1;
while (j<nSlices)
{
	setSlice(j);
label = getInfo("slice.label");
label = substring(label, 0, lengthOf(label)-2);
tCur = parseFloat(label);
diff = round((tCur-tPrev)*framerate-1);
// Find and record blank frames removed by ImageJ based on slice timestamps
for (t = 0; t < diff; t++) {
	File.append(toString(j+frameCount-1) + "." + toString(m), Deletedpath); // Add frame number to deleted - Jered S.
	m++;
	print("Blank frame recorded at time: " + toString(tPrev));
}
getRawStatistics(count, mean, min, max, std);
	if(min==255)
		{
			File.append(toString(j+frameCount-1) + "." + toString(m), Deletedpath); // Add frame number to deleted - Jered S. 
			m++;
			run("Delete Slice");
		}
		else{
		m=1; // Change deleted index to 1 - Jered S.
		j++;
		}
		tPrev = tCur;
	}
setSlice(j);
label = getInfo("slice.label");
label = substring(label, 0, lengthOf(label)-2);
tCur = parseFloat(label);
diff = round((tCur-tPrev)*framerate-1);
for (t = 0; t < diff; t++) {
	File.append(toString(j+frameCount-1) + "." + toString(m), Deletedpath); // Add frame number to deleted - Jered S.
	m++;
	print("Blank frame recorded at time: " + toString(tPrev));
}
tPrev = tCur;
getRawStatistics(count, mean, min, max, std);
if(nSlices==1 && min==255){
	print("0 Frames after deletions");
	Msg=Expname+" Finished frames: "+ toString(fend)+"/"+toString(framerange);
	print(Msg);
	continue;
}
print(toString(nSlices)+" Frames after deletions");
if (fend == 0) {
	fend = framerange;
}
frameCount=frameCount+nSlices;
//Run the tracking and save the result file
dotIndex = indexOf(Expname, ".");
Expbody = substring(Expname, 0, dotIndex)+"_"+toString(fstart)+"-"+toString(fend);
SaveDatapath=HomeDirectory+File.separator+"TkResults_"+Expbody+".txt";
call("MTrack2_.setProperty","showPathLengths","false"); 
call("MTrack2_.setProperty","showLabels","false"); 
call("MTrack2_.setProperty","showPositions","false"); 
run("MTrack2 ", "minimum=20 maximum=1000 maximum_=10000 minimum_=5 save show save=[&SaveDatapath]");
run("Close All");
Msg=Expname+" Finished frames: "+ toString(fend)+"/"+toString(framerange);
print(Msg);
}
Msg=Expname+" completed!\n";
print(Msg);
	}
}
//Print the final message
Msg="All files in the directory are completed!\n";
print(Msg);
