## Supplementary File 5 for "Evolution of Boldness and Exploratory Behavior in Giant Mice from Gough Island"

__author__ = 'Jered Stratton'
# Takes in trajectory file output from Preprocessing.ijm and calculates a variety of behavioral measures reported
# regularly at a specified time interval
# ROI: region of interest
import sys
import math
import statistics
import numpy
import calc_mobile_thresh as cmt
from collections import OrderedDict

trajFileName = sys.argv[1]  # Combined trajectory file to analyze
calibrationFileName = sys.argv[2]  # Trajectory file of calibration
cueArm = str(sys.argv[3])  # Whether cue was in left or right arm
outFilePrefix = sys.argv[4]  # Prefix for output file
outFileName = outFilePrefix + "_predatorCueData.csv"

mobileThresh = 0.0  # Distance (in pixels) needed to say a mouse is moving
mobile = False  # Indicator if subject is mobile
timeFactor = 1/30  # Number of seconds per frame
reportInterval = 60/timeFactor  # Number of frames before reporting measurements again
location = ""

positions = {'center': [0, 0], 'start': [0, 0], 'cue': [0, 0], 'control': [0, 0]}  # List of X,Y coordinates (set by calibration)
distanceFactor = 0  # Number of mm per pixel (set by calibration)
centerRadius = 0  # Radius of center ROI (in pixels, set by calibration)
armRadius = 0  # Radius of arm ROI (in pixels, set by calibration)

frameCount = 0  # Total number of frames recorded
xPrev = 0
yPrev = 0

distanceTraveled = OrderedDict([('center', 0), ('start', 0), ('cue', 0 ), ('control', 0)])  # Distance traveled in each ROI
timeMobile = OrderedDict([('center', 0), ('start', 0), ('cue', 0 ), ('control', 0)])  # Amount of time (in frames) the subject was mobile in each ROI
timeROI = OrderedDict([('center', 0), ('start', 0), ('cue', 0 ), ('control', 0)])  # Amount of time (in seconds) the subject was in each ROI
armEntrances = OrderedDict([('start', 0), ('cue', 0 ), ('control', 0)])  # Number of times the subject entered each ROI
armLatency = OrderedDict([('start', 0), ('cue', 0 ), ('control', 0)])  # Amount of time before the subject entered each arm for the first time
timePastPort = OrderedDict([('start', 0), ('cue', 0 ), ('control', 0)])  # Amount of time spent past the port in each arm (including start)
mobileLatency = 0  # Amount of time before the subject moved for the first time
distanceTraveledTotal = 0 # Total distance traveled
timeMobileTotal = 0 # Total amount of time spent mobile

output = []
# Calibrate the positions of each ROI and the distance in pixels for 1 mm
#
# Read in the calibration file and store positions in dummy objects
dummyX = []
dummyY = []
for i in range(1, 5):
    xPositions = []
    yPositions = []
    calibrationFile = open(calibrationFileName, 'r')
    calibrationFile.readline()
    calibrationFile.readline()
    for line in calibrationFile:
        line = line.rstrip()
        lineValues = line.split('\t')
        xPositions.append(float(lineValues[(i*3)-2]))
        yPositions.append(float(lineValues[(i*3)-1]))
    dummyX.append(statistics.mean(xPositions))
    dummyY.append(statistics.mean(yPositions))
    calibrationFile.close()

# Identify center
positions['center'] = [dummyX[dummyY.index(sorted(dummyY)[1])], sorted(dummyY)[1]]
print("positions of center: " + str(positions['center']))
dummyX.remove(dummyX[dummyY.index(sorted(dummyY)[1])])
dummyY.remove(sorted(dummyY)[1])
# Identify arms based on position relative to center
for i in range(0, 3):
    xCur = dummyX[i]
    yCur = dummyY[i]
    if xCur > positions['center'][0] and yCur > positions['center'][1]:
        positions['start'][0] = xCur
        positions['start'][1] = yCur
    elif xCur < positions['center'][0] and yCur > positions['center'][1]:
        if cueArm == "left":
            positions['cue'][0] = xCur
            positions['cue'][1] = yCur
        elif cueArm == "right":
            positions['control'][0] = xCur
            positions['control'][1] = yCur
        else:
            print("Arm designation not recognized")
    elif yCur < positions['center'][1]:
        if cueArm == "right":
            positions['cue'][0] = xCur
            positions['cue'][1] = yCur
        elif cueArm == "left":
            positions['control'][0] = xCur
            positions['control'][1] = yCur

# Set distanceFactor and ROI radii
centerDistances = []
for key in positions:
    distanceToCenter = math.sqrt((positions['center'][0] - positions[key][0])**2 + (positions['center'][1] - positions[key][1])**2)
    centerDistances.append(distanceToCenter)
centerDistances.remove(min(centerDistances))
print("Distances to Center:")
print(centerDistances)
distanceFactor = 539.29/statistics.mean(centerDistances)
print("Calibration at " + str(distanceFactor) + "mm per pixel")
centerRadius = 67.2/distanceFactor
armRadius = 355.6/distanceFactor

# Set Mobile threshold using method of Shoji et al.
frameCount = 0
mobileThresh, totalFrames = cmt.mobilityThreshold(trajFileName)

print("Analyzing video at " + str(1/timeFactor) + " frames per second")

# Read in trajectory file
trajFile = open(trajFileName, 'r')
trajFile.readline()
trajFile.readline()
for line in trajFile:
    line = line.rstrip()
    lineValues = line.split('\t')
    xCur = float(lineValues[1])
    yCur = float(lineValues[2])
    if frameCount == 0:
        xPrev = xCur
        yPrev = yCur
        # Check for ROI
        for key in positions:
            if key == 'center':
                if math.sqrt((positions['center'][0] - xCur)**2 + (positions['center'][1] - yCur)**2) <= centerRadius:
                    timeROI['center'] += timeFactor
                    location = "center"
            elif math.sqrt((positions[key][0] - xCur)**2 + (positions[key][1] - yCur)**2) <= armRadius:
                timeROI[key] += timeFactor
                location = key
                # Check if past port
                centerPort = math.sqrt((positions['center'][0] - positions[key][0])**2 + (positions['center'][1] - positions[key][1])**2)
                centerMouse = math.sqrt((positions['center'][0] - xCur)**2 + (positions['center'][1] - yCur)**2)
                portMouse = math.sqrt((positions[key][0] - xCur)**2 + (positions[key][1] - yCur)**2)
                if (2*centerPort*portMouse) == 0:
                    timePastPort[key] += timeFactor
                elif ((centerPort**2 + portMouse**2 - centerMouse**2)/(2*centerPort*portMouse)) <= 0:
                    timePastPort[key] += timeFactor
        frameCount += 1
        continue
    else:
        distanceCur = math.sqrt((xCur - xPrev)**2 + (yCur - yPrev)**2)
        if distanceCur > mobileThresh:
            mobile = True
            if mobileLatency == 0:
                mobileLatency = frameCount
                print(str(mobileLatency*timeFactor) + " seconds to move for the first time")
        else:
            mobile = False
        for key in positions:
            if key == 'center':
                if math.sqrt((positions['center'][0] - xCur)**2 + (positions['center'][1] - yCur)**2) <= centerRadius:
                    timeROI['center'] += timeFactor
                    location = "center"
                    if mobile:
                        distanceTraveled[key] += distanceCur * distanceFactor
                        timeMobile[key] += timeFactor
            elif math.sqrt((positions[key][0] - xCur)**2 + (positions[key][1] - yCur)**2) <= armRadius:
                timeROI[key] += timeFactor
                if location != key:
                    armEntrances[key] += 1
                location = key
                # Check if past port
                centerPort = math.sqrt((positions['center'][0] - positions[key][0]) ** 2 + (positions['center'][1] - positions[key][1]) ** 2)
                centerMouse = math.sqrt((positions['center'][0] - xCur) ** 2 + (positions['center'][1] - yCur) ** 2)
                portMouse = math.sqrt((positions[key][0] - xCur) ** 2 + (positions[key][1] - yCur) ** 2)
                if (2 * centerPort * portMouse) == 0:
                    timePastPort[key] += timeFactor
                elif ((centerPort ** 2 + portMouse ** 2 - centerMouse ** 2) / (2 * centerPort * portMouse)) <= 0:
                    timePastPort[key] += timeFactor
                if mobile:
                    distanceTraveled[key] += distanceCur * distanceFactor
                    timeMobile[key] += timeFactor
                if armLatency[key] == 0:
                    armLatency[key] = frameCount*timeFactor
                    print(str(armLatency[key]) + " seconds to enter " + str(key) + " arm for the first time")
        if mobile:
            distanceTraveledTotal += distanceCur * distanceFactor
            timeMobileTotal += timeFactor
        frameCount += 1
        if frameCount % reportInterval == 0 or frameCount == totalFrames:
            reportData = []
            reportData.append(frameCount*timeFactor)
            for key in distanceTraveled:
                reportData.append(distanceTraveled[key]/1000)
            reportData.append(distanceTraveledTotal/1000)
            for key in timeMobile:
                reportData.append(timeMobile[key])
            reportData.append(timeMobileTotal)
            for key in armEntrances:
                reportData.append(armEntrances[key])
            for key in timePastPort:
                reportData.append(timePastPort[key])
            for key in timeROI:
                reportData.append(timeROI[key])
                if timeROI[key] != 0:
                    reportData.append(distanceTraveled[key]/timeROI[key])
                else:
                    reportData.append(0)
            if timeROI['control'] > 0:
                reportData.append(timeROI['cue']/timeROI['control'])
            else:
                reportData.append("NA")
            if timeROI['start'] >0:
                reportData.append(timeROI['cue']/timeROI['start'])
                reportData.append(timeROI['control']/timeROI['start'])
            else:
                reportData.append("NA")
                reportData.append("NA")
            output.append(reportData)
        xPrev = xCur
        yPrev = yCur

trajFile.close()

outFile = open(outFileName, "w")
outFile.write("Time (seconds),Distance Traveled (m) Center,Distance Traveled (m) Start,Distance Traveled (m) Cue,"
              "Distance Traveled (m) Control,Distance Traveled (m) Total,Time Mobile Center,Time Mobile Start,"
              "Time Mobile Cue,Time Mobile Control,Time Mobile Total,Start Arm Entrances,Cue Arm Entrances,Control "
              "Arm Entrances,Time Past Start Port,Time Past Cue Port,Time Past Control Port,Time in Center,Speed "
              "in Center,Time in Start,Speed in Start,Time in Cue Arm,Speed in Cue Arm,Time in Control Arm,Speed "
              "in Control Arm,Time Cue/Control,Time Cue/Start,Time Control/Start\n")
for i in range(0, len(output)):
    outFile.write(",".join(str(x) for x in output[i]))
    outFile.write("\n")
outFile.close()
